# FMRP sustains presynaptic function via control of activity-dependent bulk endocytosis

**DOI:** 10.1101/2020.09.10.291062

**Authors:** Katherine Bonnycastle, Peter C. Kind, Michael A. Cousin

**Author notes:** Correspondence – Centre for Discovery Brain Sciences, Hugh Robson Building, George Square, University of Edinburgh, Edinburgh, Scotland, UK EH8 9XD.

## Abstract

Synaptic vesicle (SV) recycling defects are linked to neurodevelopmental disorders, including fragile X syndrome (FXS), which results from loss of fragile X mental retardation protein (FMRP) encoded by the *FMR1* gene. Hyperexcitability of neuronal circuits is a key feature of FXS, therefore we investigated whether SV recycling was affected by the absence of FMRP during increased neuronal activity. We revealed that primary neuronal cultures from a *Fmr1* knockout rat model display a specific defect in activity-dependent bulk endocytosis (ADBE). This defect resulted in an inability of *Fmr1* knockout neurons to sustain SV recycling during high frequency stimulation. Using a molecular replacement strategy, we also revealed that a human FMRP mutant that cannot bind BK channels failed to correct ADBE dysfunction in knockout neurons, however this dysfunction was corrected by BK channel agonists. Therefore, FMRP performs a key role in sustaining neurotransmitter release via selective control of ADBE, suggesting intervention via this endocytosis mode may correct hyperexcitabiltiy observed in FXS.

**SUMMARY:** Fragile X syndrome (FXS) is caused by loss of fragile X mental retardation protein (FMRP). Bonnycastle et al show that FMRP is specifically required for activity-dependent bulk endocytosis (ADBE), revealing 1) FMRP sustains neurotransmitter release and 2) intervention via ADBE may correct circuit hyperexcitabilty in FXS.

## INTRODUCTION

Information communication between neurons requires the activity-dependent, synchronous fusion of neurotransmitter-containing synaptic vesicles (SVs). SVs mobilised by action potentials (APs) are termed the recycling pool, which is subdivided into the readily releasable pool (RRP, comprising primed fusion-ready SVs) and the reserve pool (mobilised during high neuronal activity (Alabi and Tsien, 2012)).

Several SV endocytosis modes are sequentially recruited by increasing stimulus intensity to replenish these SV pools (Chanaday et al., 2019). Ultrafast endocytosis is dominant during sparse neuronal activity (Watanabe et al., 2013), whereas clathrin-mediated endocytosis becomes prevalent during AP trains (Granseth et al., 2006). Both ultrafast and clathrin-mediated endocytosis saturate during heightened neuronal activity (Lopez-Murcia et al., 2014; Soykan et al., 2017), and under these conditions, activity-dependent bulk endocytosis (ADBE) is the dominant endocytosis mode (Clayton et al., 2008). ADBE generates large endosomes directly from the plasma membrane, from which SVs are formed to replenish the reserve pool (Cheung et al., 2010; Richards et al., 2000).

The translation repressor fragile X mental retardation protein (FMRP) is located in central nerve terminals, where it controls the expression of numerous presynaptic proteins (Darnell and Klann, 2013). However, it also has protein synthesis-independent presynaptic functions, such as mediating ion channel gating, activation, and density (reviewed in Ferron, 2016). For example, FMRP interacts with Slack potassium channels (Brown et al., 2010), N-type calcium channels (Ferron et al., 2014) and large conductance voltage and calcium-gated big potassium (BK) channels (Deng et al., 2013; Deng and Klyachko, 2016; Kshatri et al., 2020). The absence of FMRP at CA3-CA1 hippocampal synapses leads to reduced BK channel activity and excessive AP broadening (Deng et al., 2013; Wang et al., 2014; Deng and Klyachko, 2016), resulting in increased presynaptic calcium influx during high frequency stimulation (Deng et al., 2011; Deng et al., 2013). These synapses also display altered short-term plasticity, but only during periods of heightened activity (Deng et al., 2011; Klemmer et al., 2011). Therefore, presynaptic phenotypes that occur in the absence of FMRP are only revealed during intense neuronal activity (Deng et al., 2011; Deng et al., 2013; Ferron et al., 2014).

Hyperexcitability of neuronal circuits is a key feature of fragile X syndrome (FXS) (Booker et al., 2019; Das Sharma et al., 2020), one of the most common monogenic causes of intellectual disability (ID) and autism spectrum disorder (Mefford et al., 2012). Most FXS cases are caused by a CGG trinucleotide expansion in the 5’ untranslated region of the *FMR1* gene which encodes FMRP, leading to hypermethylation of the promoter and silencing of the *FMR1* gene. A point mutation which disrupts polyribosome binding (I304N), is sufficient to cause FXS (Feng et al., 1997), whereas a different mutation results in mutant FMRP that is unable to bind to and regulate BK channels (R138Q).

Since FMRP may be required for accurate presynaptic function, specifically during intense neuronal activity, we determined whether SV recycling was disproportionately impacted under these conditions in primary neuronal cultures derived from a *Fmr1* knockout (KO) rat model (Asiminas et al., 2019). No significant defect in either SV exocytosis or endocytosis was observed, however *Fmr1* KO neurons displayed a robust defect in ADBE. This defect reduced presynaptic performance during periods of intense neuronal activity. Finally, molecular replacement studies with FMRP mutants revealed that the ADBE defect was due to loss of BK channel interactions and that BK channel activators could restore normal function in *Fmr1* KO neurons.

## RESULTS AND DISCUSSION

### *Fmr1* KO neurons display no obvious defect in SV recycling

FMRP is proposed to control SV fusion (Ferron et al., 2014), presynaptic AP duration (Deng et al., 2011), and short-term synaptic plasticity (Deng et al., 2011; Klemmer et al., 2011) in different murine models. To determine whether SV exocytosis or endocytosis were altered in a newly generated *Fmr1* KO rat model (Asiminas et al., 2019), we examined SV recycling using synaptophysin-pHluorin (sypHy) in primary hippocampal cultures from either KO or wild-type (WT) littermate controls. SypHy consists of the SV protein synaptophysin with a pH-sensitive EGFP (pHluorin) fused to an intraluminal loop (Granseth et al., 2006). Since sypHy reports the pH of its immediate environment, an exocytosis-dependent increase in its fluorescence signal is observed during SV fusion (Fig. 1A, B). After endocytosis, sypHy fluorescence is quenched by SV acidification. This loss of fluorescence is an estimate of the kinetics of SV endocytosis, since this is rate-limiting (Atluri and Ryan, 2006; Granseth et al., 2006). When WT neurons were challenged with a 300 AP train delivered at 10 Hz, they displayed a characteristic sypHy response, with an evoked fluorescence increase followed by an exponential post-stimulation decrease to baseline. To determine the amount of exocytosis as a proportion of the total SV pool, fluorescence traces were normalised to the sypHy response in the presence of NH_4_Cl (to reveal the maximal unquenched signal; Fig. 1B). No significant difference was observed when WT and KO were compared (Fig. 1C). The lack of effect was confirmed using the vacuolar-type ATPase inhibitor bafilomycin A1 to isolate SV exocytosis without the confound of SV endocytosis (Sankaranarayanan and Ryan, 2001) (Fig. S1A, B). The time constant, τ, of SV endocytosis was also not significantly different between WT and KO neurons (Fig. 1D). Therefore, loss of FMRP does not impair SV exocytosis or endocytosis during low frequency stimulation.

**Figure 1.**
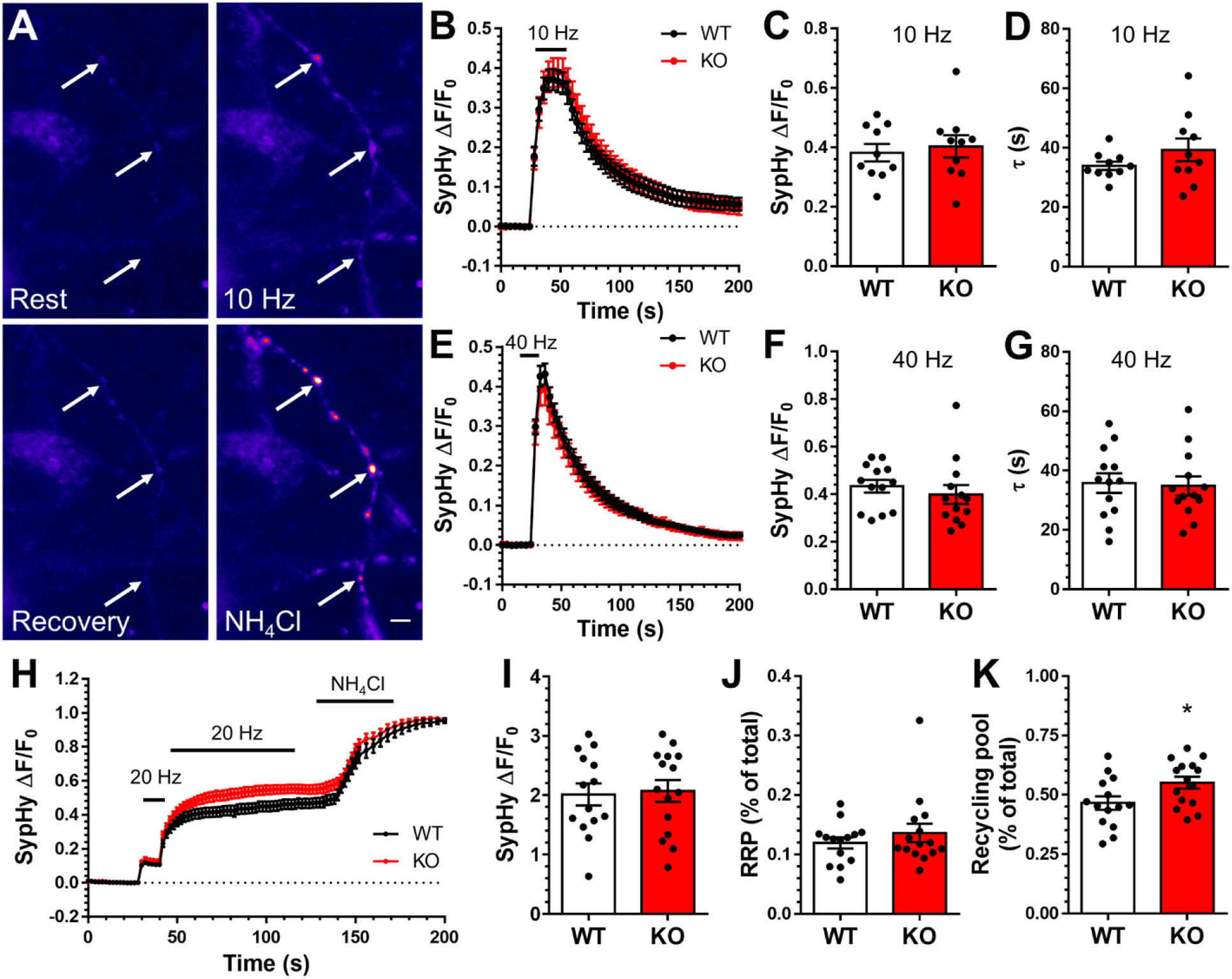
*Fmr1* KO neurons display no defect in SV recycling. Hippocampal neurons from either *Fmr1* KO or WT littermate controls were transfected with sypHy on *DIV* 7 and imaged *DIV* 13-15. A-G) Neurons were challenged with a train of APs (either 10 Hz 30 s, B-D; or 40 Hz 10 s, E-G) before exposure to NH_4_Cl 3 min later. A) Representative images of SypHy-transfected nerve terminals (indicated by arrows) during this experiment (Rest, 10 Hz, Recovery and NH_4_Cl, scale bar = 5 µm). B, E) Mean traces displaying the average sypHy response of WT (black) and KO (red) neurons in response to 10 Hz (B) or 40 Hz (E) stimulation (ΔF/F_0_ as a fraction of the total SV pool, revealed by NH_4_Cl). Bar indicates stimulation period. C, F) Mean sypHy peak heights during either 10 Hz (C) or 40 Hz (F) stimulation. D, G) Mean sypHy retrieval time constants (τ) following either 10 Hz (D) or 40 Hz (G) stimulation. H-K) SypHy transfected neurons were challenged with two AP trains (20 Hz, 2 s and 20 Hz 80 s) before exposure to NH_4_Cl in the presence of 1 μM bafilomycin A1. H) Mean traces display the average sypHy fluorescent response of WT (black) and KO (red) neurons (ΔF/F_0_, normalised to the total SV pool, revealed by NH_4_Cl). Bars indicate stimulation period. I) Mean sypHy fluorescence during exposure to NH_4_Cl. J) Mean sypHy peak height during 20 Hz 2 s stimulation (RRP, % of total pool). K) Mean sypHy peak height during 20 Hz 80 s stimulation (Recycling pool, % of total pool. B-K) All data ± SEM. For statistical information see Table S1.

Previous studies examining loss of FMRP function revealed presynaptic defects during high frequency stimulation (Deng et al., 2011; Wang et al., 2014; Ferron et al., 2014). Therefore, we next examined sypHy responses evoked by a train of 400 APs delivered at 40 Hz (Fig. 1E-G). However, there was again no difference in either the extent of SV exocytosis (Fig. 1F, Fig. S1C, D) or the kinetics of SV endocytosis (Fig. 1G). Taken together, this indicates that *Fmr1* deletion does not play a significant role in SV recycling during either low or high frequency activity.

An increase in SV pools may partially explain the observed lack of effect on SV exocytosis in *Fmr1* KO cultures. To determine this, we used bafilomycin A1 in sypHy-transfected neurons to isolate SV fusion (Fig. 1H-K). In these experiments, the RRP was mobilised by 40 APs (20 Hz), and the remainder of the recycling pool by 1600 APs (20 Hz, Fig. 1H). The resting pool (which cannot be mobilised by APs) was revealed by application of NH_4_Cl (Fig. 1H). We observed no difference either in the size of the total SV pool (Fig. 1I) or RRP between genotypes (Fig. 1J), however the SV recycling pool was increased in KO cultures (Fig. 1K). Thus, the proportions of SVs in different functional pools are shifted in KO cultures, with more SVs in the recycling pool at the expense of those in the resting pool. This adaptation suggests that *Fmr1* KO nerve terminals compensate for reduced functionality by redistributing SVs to pools that are accessible to neuronal activity.

### *Fmr1* KO neurons have defective ADBE

The increased SV recycling pool in *Fmr1* KO neurons could reflect a compensation for a presynaptic defect. Since presynaptic deficiencies in the absence of FMRP are only revealed during high frequency stimulation (Deng et al., 2011; Wang et al., 2014; Ferron et al., 2014), we next assessed whether a process that is dominant under these conditions, ADBE, was affected in *Fmr1* KO neurons. ADBE was monitored optically using uptake of 40 kDa tetramethylrhodamine (TMR)-dextran, a fluid phase marker that is selectively accumulated via this endocytosis mode (Clayton et al., 2008). WT and *Fmr1* KO neurons were challenged with a train of 400 APs (40 Hz) to maximally trigger ADBE in the presence of TMR-dextran (Clayton et al., 2008; Nicholson-Fish et al., 2015). The number of nerve terminals that perform ADBE were revealed as discrete TMR-dextran fluorescent puncta (Fig. 2A, B). We observed a significant decrease in TMR-dextran puncta in *Fmr1* KO neurons relative to WT littermate controls (Fig. 2C). Importantly, this was not due to synapse loss in the *Fmr1* KO cultures, since there was no difference in nerve terminal numbers (measured by staining for the SV protein SV2A, Fig. S2). Taken together, these results reveal a small but robust decrease in the number of nerve terminals undergoing ADBE, suggesting that FMRP may be required for this mode of endocytosis.

**Figure 2.**
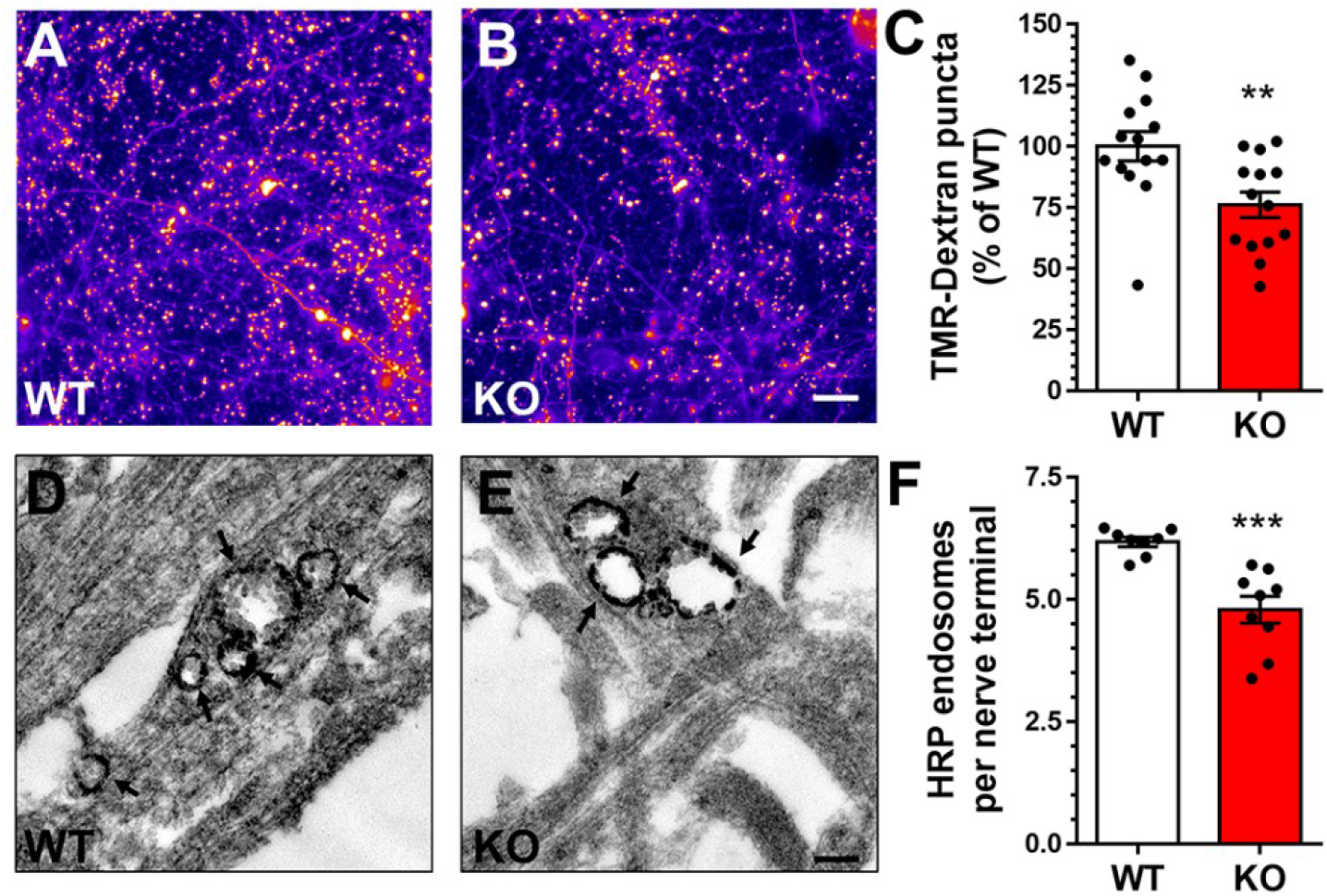
*Fmr1* KO neurons have defective ADBE. A-C) Hippocampal neurons from either *Fmr1* KO or WT littermate controls were challenged with a 40 Hz 10 s stimulus in the presence of 50 µM TMR-dextran at *DIV* 13-15. A, B) Representative images of TMR-dextran-loaded nerve terminals of WT (A) and KO (B) neurons. Scale bar = 30 µm. C) Mean TMR-dextran uptake as a proportion of total WT uptake ± SEM. D-F) *Fmr1* KO or WT neurons were challenged with a 40 Hz 10 s AP train in the presence of 10 mg/ml HRP at *DIV* 13-15. D, E) Representative images of HRP-labelled endosomes and SVs in WT (D) and KO (E) nerve terminals. Black arrows indicate HRP endosomes, scale bar = 200 nm. F) Mean HRP-labelled endosomes per nerve terminal ± SEM. For statistical information see Table S1.

TMR-dextran uptake does not report the extent to which ADBE occurs in each nerve terminal (Clayton et al., 2008). Therefore, to confirm the ADBE defect in *Fmr1* KO neurons, we applied the fluid phase marker horse radish peroxidase (HRP) during AP stimulation (40 Hz 10 s, Fig. 2D, E). Quantification of the number of HRP-labelled endosomes revealed a significant reduction in *Fmr1* KO neurons compared to WT (Fig. 2F), whereas the endosome area was not significantly different (Fig. S3). This reduction in bulk endosome generation reveals that *Fmr1* KO neurons display a specific defect in both the extent and prevalence of ADBE during intense neuronal activity.

### *Fmr1* KO neurons display decreased presynaptic performance

Since ADBE is the dominant endocytosis mode during intense neuronal activity (Clayton et al., 2008), the impact of its dysfunction on SV exocytosis may only be revealed during patterns of high frequency stimulation. To determine this, we monitored the amount of sypHy that visited the plasma membrane as a surrogate of neurotransmitter release, in response to multiple high frequency AP trains (Fig. 3A, B). We predicted that *Fmr1* KO neurons would be unable to sustain performance to the same extent as WT, due to fewer new SVs being provided via ADBE (Nicholson-Fish et al., 2015). Cultures were stimulated with four high frequency AP trains (40 Hz 10 s) separated by 5 min intervals. WT neurons displayed a sequential decrease in the extent of the evoked sypHy response, consistent with a depletion of SVs available for exocytosis. When the same protocol was performed for *Fmr1* KO neurons, the sypHy response during the final stimulation was significantly lower when compared to WT (Fig. 3B). This suggests FMRP sustains neurotransmitter release during increased neuronal activity via its control of ADBE.

**Figure 3.**
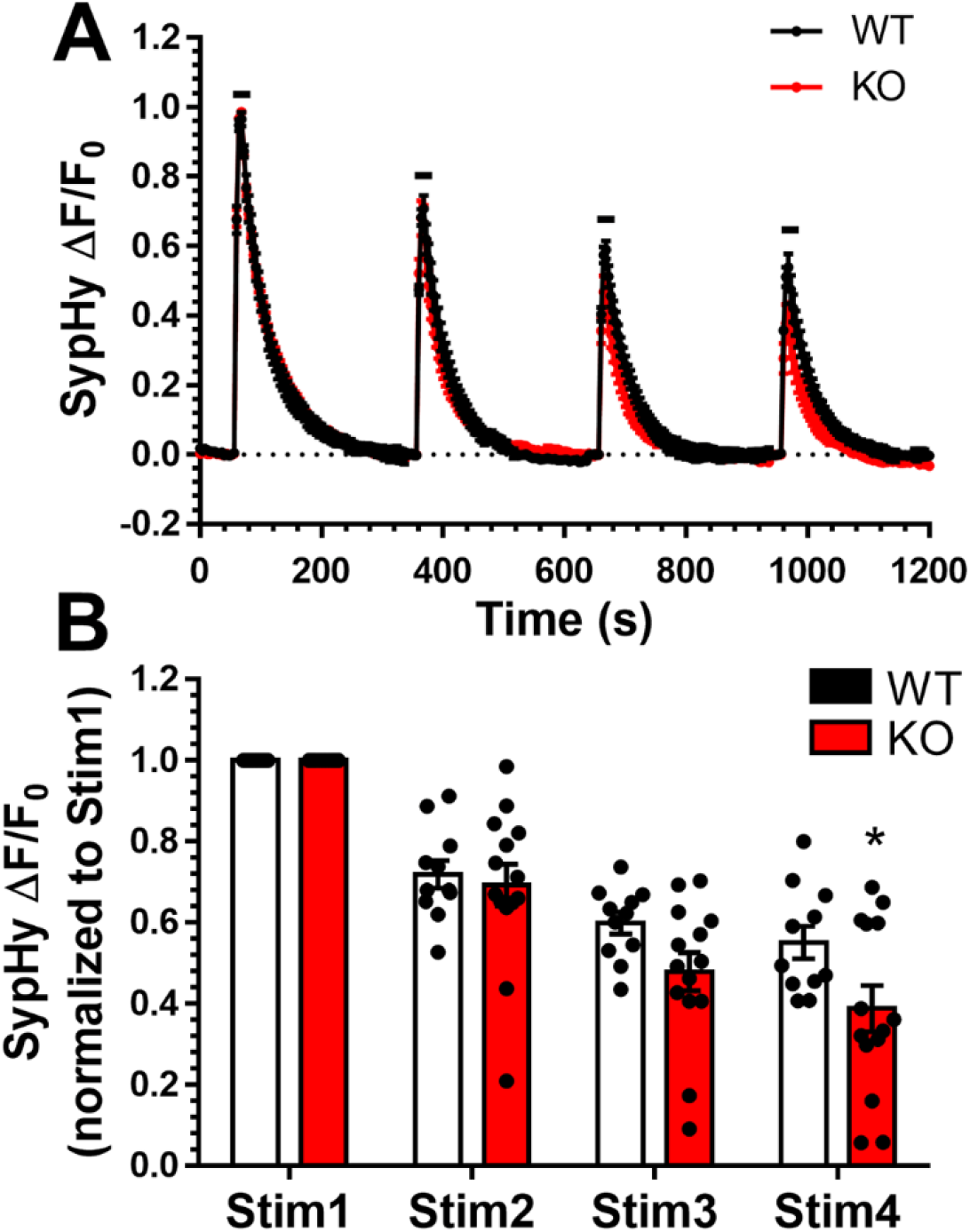
*Fmr1* KO neurons display decreased presynaptic performance. Hippocampal neurons from either *Fmr1* KO or WT littermate controls were transfected with sypHy on *DIV* 7 and imaged *DIV* 13-15. Transfected neurons were stimulated four times with 40 Hz 10 s at 5 min intervals, before exposure to NH_4_Cl. A) Mean traces displaying the average sypHy response of WT (black) and KO (red) neurons ± SEM. Traces are ΔF/F_0_ and normalised to the sypHy peak response to the first stimulus. Bar indicates stimulation period. B) Mean sypHy peak heights for each 40 Hz 10 s stimulation, normalised to the first ± SEM. For statistical information see Table S1.

### BK channel interactions are required for FMRP function in ADBE

To determine the mechanism underlying the control of ADBE by FMRP, we determined whether specific loss of function mutants could rescue TMR-dextran uptake in *Fmr1* KO cultures. First, cultures from *Fmr1* KO rats or WT littermate controls were transfected with either an empty fluorescent mCerulean vector (mCer, empty) or mCer plus EGFP-FMRP (FMRP). Overexpression of FMRP_WT_ did not affect evoked TMR-dextran uptake in WT neurons (Fig. 4A, B), indicating that excess FMRP has no detrimental effect on ADBE. It also signified that additional FMRP does not enhance ADBE, as observed with constitutively active Rab11 mutants (Kokotos et al., 2018). Importantly, FMRP_WT_ expression fully rescued the impairment in TMR-dextran uptake in *Fmr1* KO neurons (Fig. 4B).

**Figure 4.**
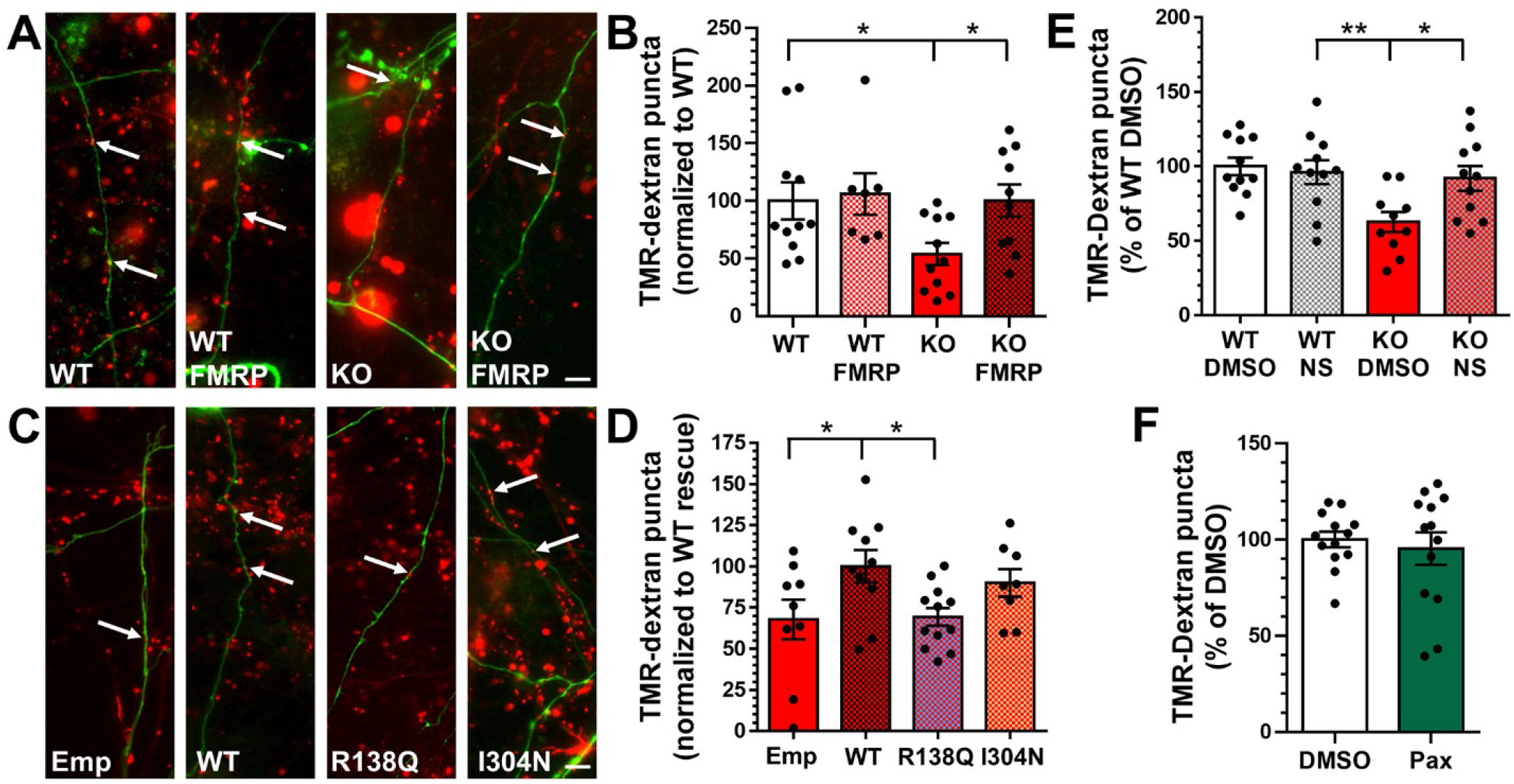
BK channel activation corrects ADBE defects in *Fmr1* KO neurons. A-D) Hippocampal neurons from either *Fmr1* KO or WT littermate controls were transfected with mCerulean (empty) or mCerulean and either GFP-FMRP_WT_ (WT), GFP-FMRP_R138Q_ (R138Q) or GFP-FMRP_I304N_ (I304N) 3 days prior to imaging. On *DIV* 13-15, neurons were challenged with a 40 Hz 10 s stimulus in the presence of 50 µM TMR-dextran. A, C) Representative images of TMR-dextran uptake (red) in axons transfected with empty or FMRP vectors (green), arrows indicate TMR-dextran puncta. Scale bar = 10 µm. B, D) Mean TMR-dextran uptake per 100 µm of transfected axon normalised to either B) WT control or D) FMRP_WT_ control (both ± SEM). E) Hippocampal neurons from either *Fmr1* KO or WT littermate controls were incubated with DMSO or 10 µM NS 11021 (NS), and 50 µM TMR-dextran for 2 min prior to a 40 Hz 10 s stimulus on *DIV* 13-15. Mean TMR-dextran uptake as a proportion of total WT uptake ± SEM. F) WT hippocampal neurons were incubated with DMSO or 10 µM Paxilline (Pax) and 50 µM TMR-dextran for 2 min prior to a 40 Hz 10 s stimulus on *DIV* 13-15. Mean TMR-dextran uptake as a proportion of total WT uptake ± SEM. For statistical information see Table S1.

We exploited the ability of FMRP_WT_ to rescue TMR-dextran uptake to perform molecular replacement studies using two loss of function mutants: FMRP_I304N_ - that does not bind ribosomes (De Boulle et al., 1993) and FMRP_R138Q_ - that does not bind BK channels (Myrick et al., 2015; Kshatri et al., 2020) (Fig. 4C). When these experiments were performed, FMRP_WT_ restored TMR-dextran uptake in *Fmr1* KO cultures as previously observed (Fig. 4D). TMR-dextran uptake in FMRP_I304N_ expressing neurons was not significantly different to neurons expressing FMRP_WT_ (Fig. 4D), indicating that FMRP-dependent control of protein translation is dispensable for ADBE. However, expression of FMRP_R138Q_ failed to restore TMR-dextran uptake in *Fmr1* KO neurons, with levels equivalent to those observed with an empty vector (Fig. 4D). These results strongly suggest that FMRP controls ADBE through interactions with BK channels.

The inability of FMRP_R138Q_ to rescue ADBE in *Fmr1* KO neurons supports the hypothesis that presynaptic defects in the absence of FMRP are translation-independent (Deng and Klyachko, 2016). The R138Q mutation is found in individuals presenting with ID (and in some cases FXS phenotypes, seizures and autism) (Myrick et al., 2015; Sitzmann et al., 2018; Diaz et al., 2018; Collins et al., 2010). However, given its prevalence in the general population (identified in 20 subjects, of which 9 were male, that display no discernable ID - https://gnomad.broadinstitute.org) it is unlikely that this mutation directly contributes to FXS.

### BK channel activation corrects ADBE defects in *Fmr1* KO neurons

The absence of ADBE rescue by FMRP_R138Q_ suggests that BK channel modulation by FMRP may be important for this endocytosis mode. Therefore, we investigated whether altering BK channel activity impacted ADBE. First, we determined whether enhancing channel activity with the activator NS 11021 could restore ADBE in *Fmr1* KO neurons. BK channel activation with NS 11021 did not affect TMR-dextran uptake in WT neurons when compared to a vehicle control (Fig. 4E). It did however restore TMR-dextran uptake in *Fmr1* KO neurons to WT levels (Fig. 4E). Therefore, activation of BK channels is sufficient to correct dysfunctional ADBE in *Fmr1* KO neurons.

To determine whether loss of BK channel function was responsible for defective ADBE in *Fmr1* KO neurons, we attempted to mimic the deficit using the BK channel antagonist Paxilline in WT hippocampal cultures. Interestingly, this manoeuvre did not impact activity-dependent TMR-dextran uptake (Fig. 4F). Taken together, these results suggest that BK channel dysfunction is not responsible for ADBE defects in *Fmr1* KO neurons, however BK channel activation can correct this fault.

### ADBE acts as a rheostat to tune neuronal excitability

It is likely that the observed reduction in ADBE reflects a compensatory or homeostatic mechanism to counteract the increased excitability in FXS and *Fmr1* KO models (Booker et al., 2019; Das Sharma et al., 2020; Zhang et al., 2014). BK channel openers decrease hyperexcitability in *Fmr1* KO models both *in vitro and in vivo* by increasing hyperpolarization, resulting in decreased calcium influx in nerve terminals.(Zhang et al., 2014; Hebert et al., 2014). The correction of ADBE by BK channel activation in this study, suggests that this is the most likely mechanism of action. Although it appears counterintuitive that an endocytosis mode which is triggered by high neuronal activity should be depressed by hyperexcitability, ADBE is reduced by stimulus frequencies above 40 Hz (Clayton et al., 2008). This inhibition may result from excess activity-dependent calcium influx, since this inhibits various forms of SV endocytosis in both large atypical and small classical central nerve terminals (Cousin and Robinson, 2000; Wu and Wu, 2014; von Gersdorff and Matthews, 1994; Leitz and Kavalali, 2016).

A homeostatic reduction in ADBE in response to increased hyperexcitability would not be a homogenous adaptation across the brain, since neurons that display high firing rates would be disproportionately impacted. Therefore, even if reduced ADBE is not a causal mechanism in FXS, it may still be a valuable therapeutic intervention point to correct hyperexcitability in specific circuits (Booker et al., 2019; Das Sharma et al., 2020). Future studies are now required to determine whether modulation of ADBE can sculpt circuit activity in FXS (and other autism models that display hyperexcitability - such as SynGAP haploinsufficiency disorder (Gamache et al., 2020)). However, irrespective of whether ADBE dysfunction directly contributes to FXS, the role of FMRP in the control of this key event provides valuable new information into the mechanisms of SV recycling that regulate presynaptic function.

## MATERIALS AND METHODS

### Materials

Unless otherwise specified, all cell culture reagents were obtained from ThermoFisher Scientific (Paisley, UK). Foetal bovine serum was from Biosera (Nuaille, France). Papain was obtained from Worthington Biochemical (Lakewood, NJ, USA). All other reagents were obtained from Sigma-Aldrich (Poole, UK). Rabbit anti-SV2A was obtained from Abcam (Cambridge, UK; ab32942; RRID: AB_778192). Anti-rabbit Alexa Fluor 488 was obtained from Invitrogen (Paisley, UK; A11008; RRID AB_143165).

### Animals

Animal work was performed in accordance with the UK Animal (Scientific Procedures) Act 1986, under Project and Personal Licence authority and was approved by the Animal Welfare and Ethical Review Body at the University of Edinburgh (Home Office project licence – 7008878). Specifically, all animals were killed by schedule 1 procedures in accordance with UK Home Office Guidelines; adults were killed by exposure to CO_2_ followed by decapitation, whereas embryos were killed by decapitation followed by destruction of the brain. Heterozygous LE-*Fmr1*^em1/PWC^ female rats (Asiminas et al., 2019) were mated with WT males to produce either *Fmr1* KO or WT males. Male embryos were taken at e18.5-e19.5 for hippocampal dissection.

### DNA constructs

Site-directed mutagenesis was used to introduce single base pair mutations into the hEGFP-FMRP isoform 1 plasmid (Khayachi et al., 2018) obtained from Dr B. Bardoni (INSERM, IPMC CNRS). hEGFP-FMRP_R138Q_ and hEGFP-FMRP_I304N_ were generated using site-targeted mutagenesis (R138Q forward primer GTGCCAGAAGACTTAC<u>A</u>GCAAATGTGTGCCAAA; reverse primer TTTGGCACACATTTGC<u>T</u>GTAAGTCTTCTGGCAC: I304N forward primer AAAAATGGAAAGCTGA<u>A</u>TCAGGAGATTGTGGAC; reverse GTCCACAATCTCCTGA<u>T</u>TCAGCTTTCCATTTTT (mutated bases underlined). The base change was confirmed by Source Bioscience Sanger Sequencing (Glasgow, UK). The mCerN1 (empty vector) was constructed as previously described (Cheung et al., 2010). Synaptophysin-pHluorin (sypHy) (Granseth et al., 2006) was provided by Prof. L. Lagnado (University of Sussex, UK).

### Primary hippocampal neuronal cultures and transfection

Hippocampi from each embryo were processed separately to avoid contamination across genotypes. Dissociated primary hippocampal cultures were prepared from embryos as previously described (Zhang et al., 2015). Briefly, isolated hippocampi were digested in a 10 U/mL papain solution (Worthington Biochemical, LK003178) at 37°C for 20 min. The papain was then neutralised using DMEM F12 (ThermoFisher Scientific, 21331-020) supplemented with 10 % Foetal bovine serum (BioSera, S1810-500) and 1 % penicillin/streptomycin (ThermoFisher Scientific, 15140-122). Cells were triturated to form a single cell suspension and plated at 5 × 10^4^ cells (with the exception of single cell TMR-dextran uptake experiments, 2.5 × 10^4^ cells) per coverslip on laminin (10 µg/ mL; Sigma Aldrich, L2020) and poly-D-lysine (Sigma Aldrich, P7886) coated 25 mm glass coverslips (VWR International Ltd, Lutterworth, UK). Cultures were maintained in Neurobasal media (ThermoFisher Scientific, 21103-049) supplemented with 2 % B-27 (ThermoFisher Scientific, 17504-044), 0.5 mM L-glutamine (ThermoFisher Scientific, 25030-024) and 1% penicillin/streptomycin. After 2-3 days *in vitro* (DIV), 1 µM of cytosine arabinofuranoside (Sigma Aldrich, C1768) was added to each well to inhibit glial proliferation. Hippocampal neurons were transfected with sypHy at *DIV* 7 using Lipofectamine 2000 (ThermoFisher Scientific, 11668027) as per manufacturer’s instructions and imaged at *DIV* 13-15. For single-cell dextran experiments, neurons were transfected with mCerN1 or hEGFP-FMRP constructs using Lipofectamine 2000 3 days prior to imaging at *DIV* 13-15.

### Fluorescence imaging of sypHy

SypHy-transfected neurons were visualised at 500 nm band pass excitation with a 515 nm dichroic filter and a long-pass >520 nm emission filter on a Zeiss Axio Observer D1 inverted epifluorescence microscope (Cambridge, UK). Images were captured using an AxioCam 506 mono camera (Zeiss) with a Zeiss EC Plan Neofluar 40x/1.30 oil immersion objective. Image acquisition was performed using Zen Pro software (Zeiss). Hippocampal cultures were mounted in a Warner Instruments (Hamden, CT, USA) imaging chamber with embedded parallel platinum wires (RC-21BRFS) and challenged with field stimulation using a Digitimer LTD MultiStim system-D330 stimulator (current output 100 mA, current width 1 ms) either at 10 Hz for 30 s, 40 Hz for 10 s or 4X 40 Hz for 10 s with 5 min intervals between trains. Imaging time courses were acquired at 4 s intervals while undergoing constant perfusion with imaging buffer (119 mM NaCl, 2.5 mM KCl, 2 mM CaCl_2_, 2 mM MgCl_2_, 25 mM HEPES, 30 mM glucose at pH 7.4, supplemented with 10 μM 6-cyano-7-nitroquinoxaline-2,3-dione (Abcam, Cambridge, UK, ab120271) and 50 μM DL-2-Amino-5-phosphonopentanoic acid (Abcam, Cambridge, UK, ab120044). NH_4_Cl alkaline buffer (50 mM NH_4_Cl substituted for 50 mM NaCl) was used to reveal the maximal pHluorin response.

### Measurement of functional SV pool size

The size of the different functional SV pools in hippocampal neurons were measured by stimulating sypHy-transfected neurons for increasing durations in the presence of 1 µM bafilomycin A1 (Cayman Chemical Company, Ann Arbor Michigan, USA, 11038). The RRP was mobilised by 40 APs (20 Hz), and 10 s later the remainder of the recycling pool mobilised with a second challenge of 1600 APs (20 Hz). The resting pool was revealed by application of NH_4_Cl buffer.

### Analysis of sypHy fluorescence traces

Time traces were analysed using the FIJI distribution of Image J (National Institutes of Health). Images were aligned using the Rigid body model of the StackReg plugin (https://imagej.net/StackReg) (Thevenaz et al., 1998). Nerve terminal fluorescence was measured using the Time Series Analyser plugin (https://imagej.nih.gov/ij/plugins/time-series.html) (Balaji and Ryan, 2007). Regions of interest (ROIs) 5 pixels in diameter were placed over nerve terminals that responded to the electrical stimulus. A response trace was calculated for each cell by averaging the individual traces from each selected ROI. Fluorescence decay time constants (tau, τ, s) were calculated by fitting a monoexponential decay curve to data from the time point after the end of electrical stimulation.

### Fluorescence imaging of TMR-dextran uptake

TMR-dextran (ThermoFisher Scientific, D1842) uptake was performed as described previously (Nicholson-Fish et al., 2015). Neurons were mounted on a Zeiss Axio Observer D1 microscope as described above before challenging with 400 action potentials (40 Hz) in the presence of 50 µM of TMR-dextran (40,000 MW) in imaging buffer. Where the experiment was performed with BK channel modulators, neurons were incubated in 50 µM of TMR-dextran and either DMSO (Sigma Aldrich, D8418), 10 µM NS 11021 (Bio-Techne Ltd, Abingdon, UK, 4788/10) or 10 µM Paxilline (Bio-Techne Ltd, Abingdon, UK, 2006/10) for 120 s prior to stimulation. The TMR-dextran solution was immediately washed away after stimulation terminated, and images were acquired using 556/25 nm excitation and 630/98 nm emission bandpass filters (Zeiss) while undergoing constant perfusion. Per coverslip of cells, 3-6 different fields of view were imaged. The TMR-dextran puncta in each image were quantified using the Analyze Particles plugin of Image J (NIH, https://imagej.nih.gov/ij/developer/api/ij/plugin/filter/ParticleAnalyzer.html) to select and count particles of 0.23-0.91 µm^2^.

Where TMR-dextran uptake was performed on transfected cultures, WT and KO neurons were transfected 3 days prior to imaging with mCer-N1 plus either hEGFP-FMRP_WT_, hEGFP-FMRP_I304N_ or hEGFP-FMRP_R138Q_ or mCer-N1 alone. Transfected axons were visualised on *DIV* 13-15 at both 430 nm and 500 nm excitation (long-pass emission filter >520 nm) to ensure co-transfection. Images of transfected neurons and of TMR-dextran were acquired using 470/27 nm and 556/25 nm double band pass filters and emission filters 512/30 nm and 630/98 nm respectively (Zeiss). Per coverslip of cells, 2-8 neurons were imaged. Axon length was calculated using the Simple Neurite Tracer plugin of Image J (NIH, https://imagej.net/SNT). TMR-dextran puncta (0.23-0.91 µm^2^) were counted along transfected axons with the final value normalised for axon length. For all experiments, for each condition, at least one unstimulated coverslip was imaged to correct for the background level of TMR-dextran uptake.

### Immunofluorescence staining

Immunofluorescence staining was performed as previously described (Nicholson-Fish et al., 2015). Briefly, hippocampal neurons were fixed with 4 % paraformaldehyde (Sigma Aldrich, 47608) in PBS for 15 min. Excess paraformaldehyde was quenched with 50 mM NH_4_Cl in PBS. Cells were then permeabilized in 1 % bovine serum albumin (BSA; Roche Diagnostics GmbH, Germany, 10735078001) in PBS-Triton 0.1 % solution for 5 min and blocked in 1 % BSA in PBS at room temperature for 1 h. After blocking, cells were incubated in rabbit anti-SV2A (1:200 dilution) for 1 h, after which the cultures were washed with PBS and incubated with fluorescently conjugated secondary antibodies (anti-rabbit Alexa Fluor 488; 1:1000 dilution) for 1 hr. The coverslips were mounted on slides for imaging with FluorSave (Millipore, Darmstadt, Germany, 345789). SV2A puncta were visualised at 500 nm band pass excitation with a 515 nm dichroic filter and a long-pass >520 nm emission filter on a Zeiss Axio Observer D1 inverted epifluorescence microscope (Cambridge, UK). Images were captured using an AxioCam 506 mono camera (Zeiss) with a Zeiss EC Plan Neofluar 40x/1.30 oil immersion objective. SV2A puncta in each image were quantified using the Analyze Particles plugin of Image J to select and count particles of 0.23-3.18 µm^2^.

### HRP uptake

Hippocampal cultures were mounted in the RC-21BRFS stimulation chamber and challenged with 400 action potentials (40 Hz) in the presence of 10 mg/ml HRP (Sigma Aldrich, P8250) supplemented imaging buffer. Immediately following the end of stimulation, cultures were washed in imaging buffer to remove non-internalised HRP and fixed with a solution of 2 % glutaraldehyde (Electron Microscopy Sciences, Hatfield, USA, 16019) in phosphate buffered saline. After washing in 0.1 M Tris buffer, HRP was developed with 0.1 % 3,3’-diaminobenzidine (Fluka Chemica, Gillingham, UK, 22204001) and 0.2 % v/v hydrogen peroxide (Honeywell, Muskegon, USA, 216763) in Tris buffer. After further washing in Tris buffer, cultures were then stained with 1 % osmium tetroxide (TAAB laboratory and microscopy, Aldermaston, UK, O015/1) for 30 min. Samples were then dehydrated using an ethanol series and polypropylene oxide (Electron Microscopy Sciences, Hatfield, USA, 20411) and embedded using Durcupan resin (Sigma Aldrich, 44610). Samples were sectioned, mounted on grids, and viewed using an FEI Tecnai 12 transmission electron microscope (Oregon, USA). Intracellular structures that were <61 nm in diameter were arbitrarily designated to be SVs, whereas larger structures were considered endosomes. The endosome area was obtained by tracing the circumference using the freehand selections tool in ImageJ and measuring the resulting area. Typically, 30 fields of view were acquired for one coverslip of cells. The average number of HRP-labelled endosomes and SVs per nerve terminal was calculated for each coverslip and represents the experimental n.

### Quantification and data analysis

Microsoft Excel (Microsoft, Washington, USA) and Prism 8 software (GraphPad software Inc., San Diego USA) were used for data processing and analysis. The experimenter was blinded during both the acquisition and analysis of data. For all figures, results are presented with error bars as ± SEM, and the n for each condition represents the number of coverslips imaged. For all assays, cells were obtained from at least two cultures (N), with independent preparations from at least three individual embryos.

## ACKNOWLEDGEMENTS

Work is supported by the Simons Foundation (529508), The RS McDonald fund and a College of Medicine and Veterinary Medicine studentship. We thank Jennifer Darnell for expert advice and Steven Mitchell for excellent technical assistance. The authors declare no competing financial interests.

## AUTHOR CONTRIBUTIONS

Conceptualization, PCK, MAC; Methodology, KB, MAC; Data analysis, KB; Investigation, KB, MAC; Resources, PCK; Writing – All; Funding Acquisition, PCK, MAC.

## ABBREVIATIONS

SV: synaptic vesicle
FMRP: fragile X mental retardation protein
FXS: fragile X syndrome
AP: action potential
ADBE: activity-dependent bulk endocytosis
BK: big potassium
RRP: readily releasable pool
ID: intellectual disability
HRP: horseradish peroxidase
KO: knockout
WT: wild-type
DIV: day *in vitro*
TMR: tetramethylrhodamine
BSA: bovine serum albumin
ROI: region of interest
mCer: mCerulean
sypHy: synaptophysin-pHluorin

---

**Figure S1.**
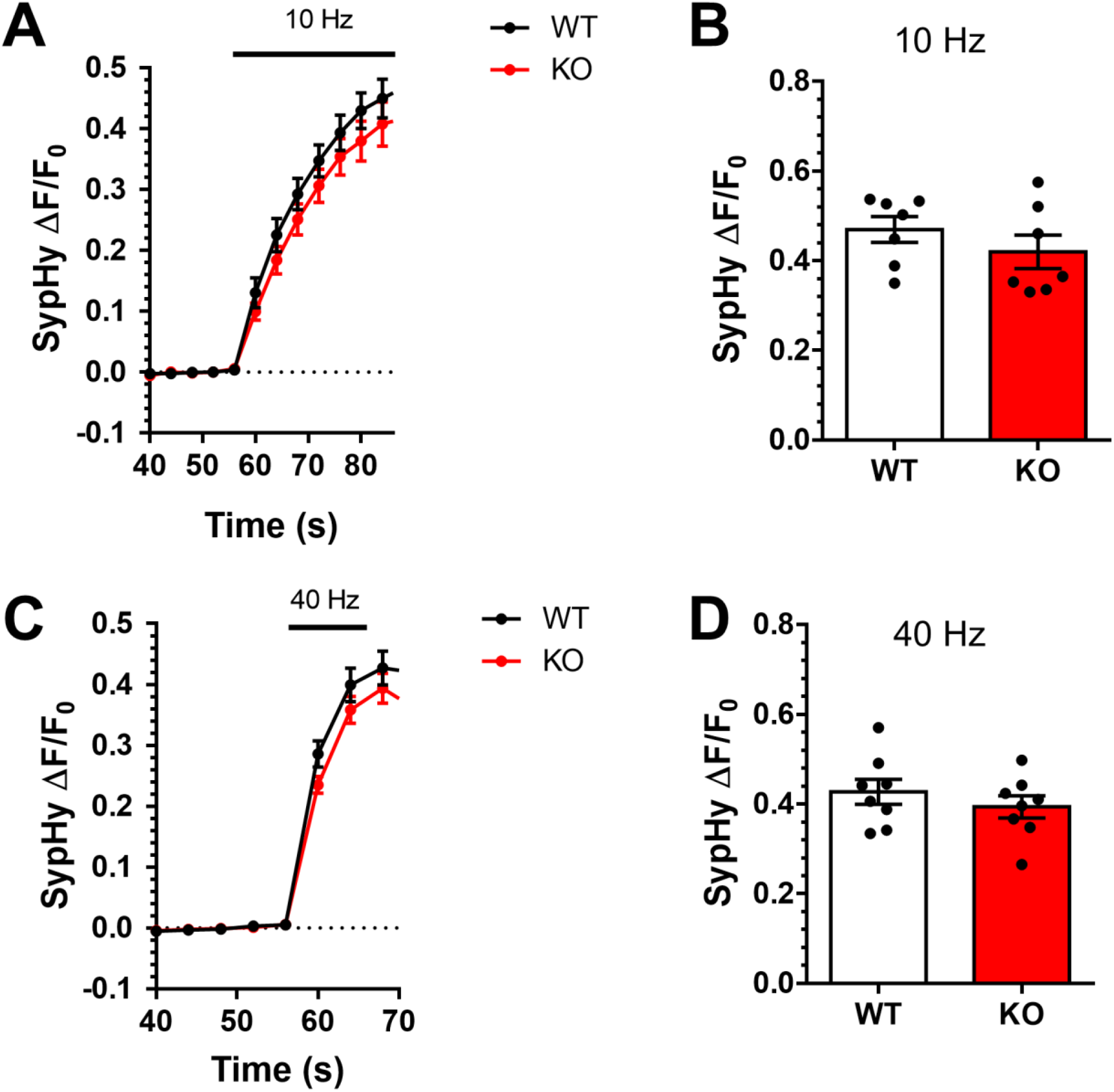
No difference in SV exocytosis across genotypes. Hippocampal neurons from either *Fmr1* KO or WT littermate controls were transfected with sypHy on *DIV* 7 and imaged *DIV* 13-15 in the presence of 1 μM bafilomycin A1. Neurons were challenged with a train of APs (either 10 Hz 30 s, A-B; or 40 Hz 10 s, C-D) before exposure to NH_4_Cl. A, C) Mean traces displaying the average sypHy response of WT (black) and KO (red) neurons in response to 10 Hz (A) or 40 Hz (C) stimulation (ΔF/F_0_ as a fraction of the total SV pool, revealed by NH_4_Cl). Bar indicates stimulation period. B, D) Mean sypHy peak heights during either 10 Hz (B) or 40 Hz (D) stimulation. A-D) All data shown ± SEM. For statistical information see Table S1.

**Figure S2.**
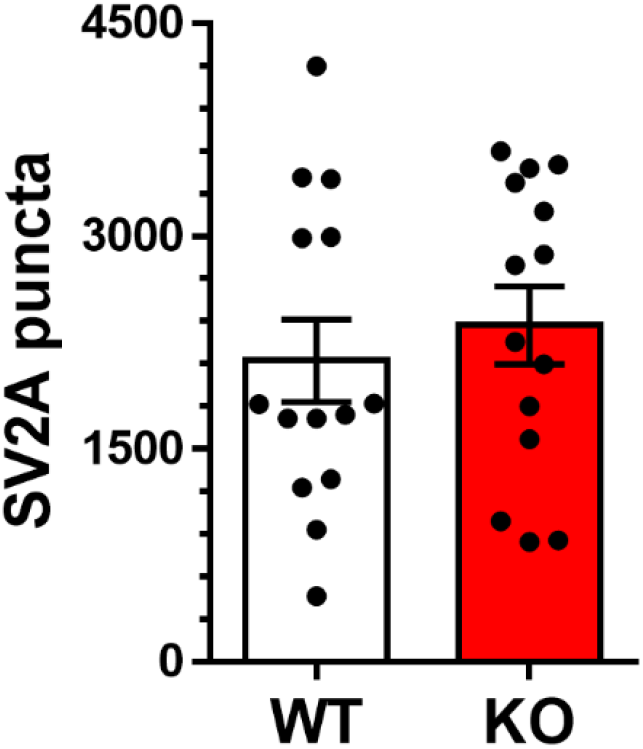
No difference in nerve terminal number between genotypes. Hippocampal neurons derived from either *Fmr1* KO or WT littermate controls were fixed and stained with anti-SV2A as a marker of nerve terminals. Bar graph represents the mean number of SV2A-stained puncta for both KO and WT ± SEM, For statistical information see Table S1.

**Figure S3.**
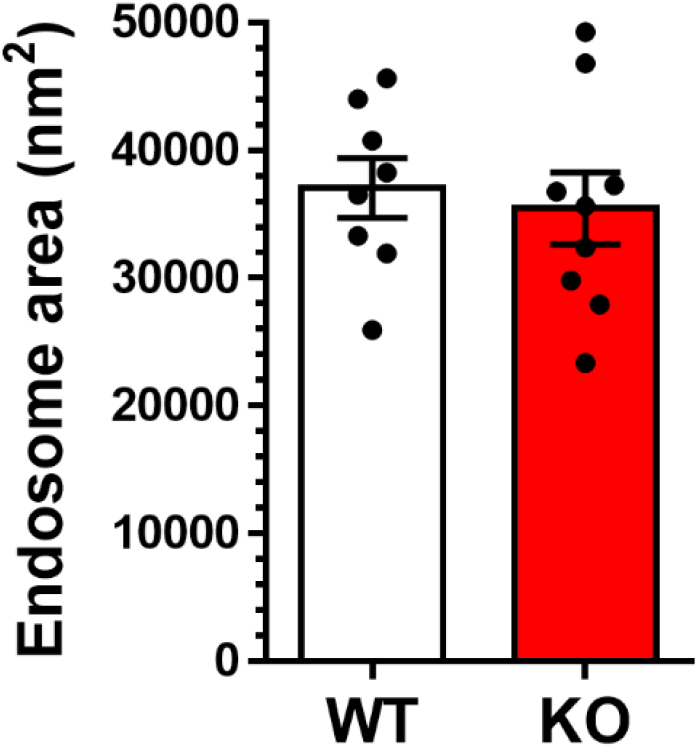
No difference in endosome size across genotypes. Hippocampal neurons derived from either *Fmr1* KO or WT littermate controls were challenged with a 40 Hz 10 s action potential train in the presence of 10 mg/ml HRP at *DIV* 13-15. Mean HRP-labelled endosome areas in WT and KO ± SEM. For statistical information see Table S1.

**Table S1.** Statistical information.

| Figure 1 | Group | Mean±SEM | n = # of coverslips/ N = # of neuronal preparations | Comparison | P | Statistical test |
| --- | --- | --- | --- | --- | --- | --- |
| Fig1C | WT | 38.2 ± 2.9 % | 10/4 | WT vs KO | 0.6545 | Unpaired t-test |
|  | KO | 40.4 ± 3.7 % | 10/4 |  |  |  |
| Fig1D | WT | 34.0 ± 1.4 s | 10/4 | WT vs KO | 0.2076 | Unpaired t-test |
|  | KO | 39.3 ± 3.8 s | 10/4 |  |  |  |
| Fig1F | WT | 43.3 ± 2.6 % | 13/4 | WT vs KO | 0.4751 | Unpaired t-test |
|  | KO | 39.9 ± 3.9 % | 13/4 |  |  |  |
| Fig1G | WT | 31.8 ± 2.9 s | 13/4 | WT vs KO | 0.8382 | Unpaired t-test |
|  | KO | 31.1 ± 2.8 s | 13/4 |  |  |  |
| Fig1I | WT | 2.0 ± 0.2 u | 14/3 | WT vs KO | 0.8275 | Unpaired t-test |
|  | KO | 2.0 ± 0.2 u | 15/3 |  |  |  |
| Fig1J | WT | 12.0 ± 0.9 % | 14/3 | WT vs KO | 0.3715 | Unpaired t-test |
|  | KO | 13.6 ± 1.5 % | 15/3 |  |  |  |
| Fig1K | WT | 46.5 ± 2.8 % | 14/3 | WT vs KO | 0.0299 | Unpaired t-test |
|  | KO | 55.2 ± 2.5 % | 15/3 |  |  |  |
| Figure 2 | Group | Mean±SEM | n = # of coverslips/ N = # of neuronal preparations | Comparison | P | Statistical test |
| Fig2C | WT | 100.0 ± 16.2 % | 14/3 | WT vs KO | 0.0054 | Unpaired t-test |
|  | KO | 76.0 ± 5.2 % | 14/3 |  |  |  |
| Fig2F | WT | 6.2 ± 0.1 endosomes | 8/3 | WT vs KO | 0.0004 | Unpaired t-test |
|  | KO | 4.8 ± 0.3 endosomes | 9/3 |  |  |  |
| Figure 3 | Group | Mean±SEM | n = # of cells or coverslips/ N = # of neuronal preparations | Comparison | P | Statistical test |
| Fig3B | WT Stim1 | 1.00 ± 0.00 u | n 11/2 | WT Stim1 vs KO Stim1 | >0.9999 | 2-way ANOVA with Bonferroni's multiple comparison test |
|  | KO Stim1 | 1.00 ± 0.00 u | n 14/2 |  |  |  |
|  | WT Stim2 | 0.72 ± 0.03 u | n 11/2 | WT Stim2 vs KO Stim2 | >0.9999 |  |
|  | KO Stim2 | 0.69 ± 0.05 u | n 14/2 |  |  |  |
|  | WT Stim3 | 0.60 ± 0.03 u | n 11/2 | WT Stim3 vs KO Stim3 | 0.1529 |  |
|  | KO Stim3 | 0.48 ± 0.05 u | n 14/2 |  |  |  |
|  | WT Stim4 | 0.55 ± 0.04 u | n 11/2 | WT Stim4 vs KO Stim4 | 0.0217 |  |
|  | KO Stim4 | 0.39 ± 0.06 u | n 14/2 |  |  |  |

Table S1. **Statistical information**
| Figure 4 | Group | Mean±SEM | n = # of coverslips/ N = # of neuronal preparations | Comparison | P | Statistical test |
| --- | --- | --- | --- | --- | --- | --- |
| Fig4B | WT | 100.0 ± 16.2 % | 11/4 | WT vs KO | 0.043 | 1-way ANOVA with Bonferroni's multiple comparisons test |
|  | WT FMRP | 106.1 ± 18.1 % | 7/4 |  |  |  |
|  | KO | 54.1 ± 9.6 % | 11/4 | KO vs KO FMRP | 0.048 |  |
|  | KO FMRP | 100.2 ± 14.0 % | 10/4 |  |  |  |
| Fig4D | Empty (Emp) | 67.8 ± 12.1 % | 8/3 | WT vs empty | 0.043 | 1-way ANOVA with Dunnett's multiple comparison test |
|  | WT | 100.0 ± 9.8 % | 10/3 | WT vs R138Q | 0.038 |  |
|  | R138Q | 69.4 ± 5.4 % | 8/3 | WT vs I304N | 0.788 |  |
|  | I304N | 89.9 ± 8.4 % | 12/3 |  |  |  |
| Fig4E | WT DMSO | 100.0 ± 5.9 % | 11/3 | WT DMSO vs KO DMSO | 0.0039 | 1-way ANOVA with Bonferroni's multiple comparisons test |
|  | WT NS | 96.1 ± 8.0 % | 11/3 | KO DMSO vs KO NS | 0.0322 |  |
|  | KO DMSO | 62.6 ± 6.8 % | 10/3 | WT DMSO vs KO NS | >0.9999 |  |
|  | KO NS | 91.9 ± 8.3 % | 11/3 | WT DMSO vs WT NS | >0.9999 |  |
| Fig4F | DMSO | 100.0 ± 4.1 % | 13/3 | DMSO vs Paxilline | 0.6182 | Unpaired t-test |
|  | Paxilline | 95.3 ± 8.4 % | 13/3 |  |  |  |
| Figure S1 | Group | Mean±SEM | n = # of coverslips/ N = # of neuronal preparations | Comparison | P | Statistical test |
| FigS1B | WT | 46.9 ± 2.8 % | 7/3 | WT vs KO | 0.3123 | Unpaired t-test |
|  | KO | 42.0 ± 3.7 % | 7/3 |  |  |  |
| FigS1D | WT | 42.7 ± 2.8 % | 8/2 | WT vs KO | 0.3809 | Unpaired t-test |
|  | KO | 39.4 ± 2.4 % | 8/2 |  |  |  |
| Figure S2 | Group | Mean±SEM | n = # of coverslips/ N = # of neuronal preparations | Comparison | P | Statistical test |
| FigS2A | WT | 2123.0 ± 291.3 puncta | 14/3 | WT vs KO | 0.5382 | Unpaired t-test |
|  | KO | 2372.0 ± 274.1 puncta | 14/3 |  |  |  |
| Figure S3 | Group | Mean±SEM | n = # coverslips/ N = # of neuronal preparation, | Comparison | P | Statistical test |
| FigS3A | WT | 37056 ± 2322 nm | n 8/3 | WT vs KO | 0.6743 | Two-way ANOVA |
|  | KO | 35468 ± 2812 nm | n 9/3 |  |  |  |

